# Corollary Discharge versus Efference Copy: Distinct Neural Signals in Speech Preparation Differentially Modulate Auditory Responses

**DOI:** 10.1101/2020.01.14.905620

**Authors:** Siqi Li, Hao Zhu, Xing Tian

## Abstract

Actions influence sensory processing in a complex way to shape behavior. For example, during actions, a copy of motor signals—termed *corollary discharge* (*CD*) or *efference copy* (*EC*)—can be transmitted to sensory regions and modulate perception. However, the sole inhibitory function of the motor copies is challenged by mixed empirical observations as well as multifaceted computational demands for behaviors. We hypothesized that the content in the motor signals available at distinct stages of actions determined the nature of signals (*CD* vs. *EC*) and constrained their modulatory functions on perceptual processing. We tested this hypothesis using speech in which we could precisely control and quantify the course of action. In three electroencephalography (EEG) experiments using a novel delayed articulation paradigm, we found that preparation without linguistic contents suppressed auditory responses to all speech sounds, whereas preparing to speak a syllable selectively enhanced the auditory responses to the prepared syllable. A computational model demonstrated that a bifurcation of motor signals could be a potential algorithm and neural implementation to achieve the distinct functions in the motor-to-sensory transformation. These results suggest that distinct motor signals are generated in the motor-to-sensory transformation and integrated with sensory input to modulate perception.

## Introduction

Actions influence sensory processing in a complex way to shape behavior. For example, the theory of internal forward models (Kawato 1999; Wolpert and Ghahramani 2000; Schubotz 2007) proposes that during actions, a copy of motor signals, independently coined as *corollary discharge* (*CD*) by Sperry (Sperry 1950) and *efference copy* (*EC*) by von Holst and Mittelstaedt (von Holst and Mittelstaedt 1950), can be transmitted to sensory regions and serves as a predictive signal to modulate sensory processing and perception. The common presumption regarding the functions of *CD* and *EC* is sensory suppression -- these motor signals suppress sensory processing via motor-to-sensory transformation in given modalities. Based on the inhibitory functions, various cognitive abilities and behaviors can be achieved, such as efficient motor control (Miall and Wolpert 1996; Kawato 1999; Wolpert and Ghahramani 2000), stable visual perception (Ross et al. 2001; Sommer and Wurtz 2006), fluent vocal and speech production and control (Guenther 1995; Tian and Poeppel 2010; Houde and Nagarajan 2011; Hickok 2012), self-monitoring and agency (Blakemore and Decety 2001; Grush 2004; Desmurget et al. 2009). Such motor-to-sensory transformation mechanisms have been evident among animal species (Crapse and Sommer 2008), and their neural pathways have been increasingly mapped out (Poulet and Hedwig 2006; Schneider et al. 2014, 2018).

However, the advances in anatomical and functional evidence for the motor-to-sensory transformation bring discrepancies. Computationally, On the one hand, the stability of perception during visual saccade requires motor signals to *suppress* the processing of sensory feedback (Ross et al. 2001). On the other hand, the predictive nature of motor signals mediates the receptive field remapping and *enhances* the sensory and perceptual sensitivity (Mohr et al. 2003; Neuweiler 2003). Empirically, in addition to the commonly observed action-induced-suppression in the sensory systems, action-induced-enhancement has also been found in a subset of sensory cortices. (Eliades and Wang 2005, 2008; Flinker et al. 2010; Singla et al. 2017; Enikolopov et al. 2018). Cognitively, higher-level cognitive functions, such as self-monitoring, require motor signals to suppress the feedback to indicate the consequences of self-induced actions (Blakemore and Decety 2001; Grush 2004; Desmurget et al. 2009). Whereas for working memory and mental imagery, positive neural representations are needed to establish the mental images (Tian and Poeppel 2010, 2012, 2013; Mary Zarate et al. 2015; Tian et al. 2016, 2018; Ma and Tian 2019). Clinically, the misattribution of inner speech to external sources in psychosis has been linked to the malfunction of agency via broken inhibitory functions from the motor system (Ford and Mathalon 2004). However, the positive symptoms, such as auditory hallucinations, require activating specific perceptual representations without corresponding external stimulations (Waters et al. 2012). The mixed observations and competing functions in motor-to-sensory transformation necessitate a reconsideration of the theoretical framework.

The observed action-induced sensory modulation occurs mostly during or after the execution of actions (Blakemore et al. 1998; Yang et al. 2008; Aliu et al. 2009; Schneider et al. 2014; Cavanaugh et al. 2016), with a few studies indicating the occurrence immediately before the execution (Eliades and Wang 2003; Daliri and Max 2016). Arguably, the execution phase is the last stage along with the entire action dynamics that include at least the intention and preparation stages (Fig. 1). These early stages are mediated by the upper-stream motor circuitries that could potentially provide different motor signals (Crapse and Sommer 2008; Straka et al. 2018). The last execution stage could bundle all the available motor signals and yield the observed mixed results and competing functions. We hypothesize, similar to previous theoretical proposals (Crapse and Sommer 2008; Straka et al. 2018), that copies of distinct motor signals are available to transmit to sensory regions at distinct temporal stages during actions (Fig. 1). Specifically, the *CD* is available right after the initiation of action dynamics -- the intention of movement, whereas the *EC* is available only after the development of a concrete movement plan -- the encoding of movement (Fig. 1).

**Figure 1.**
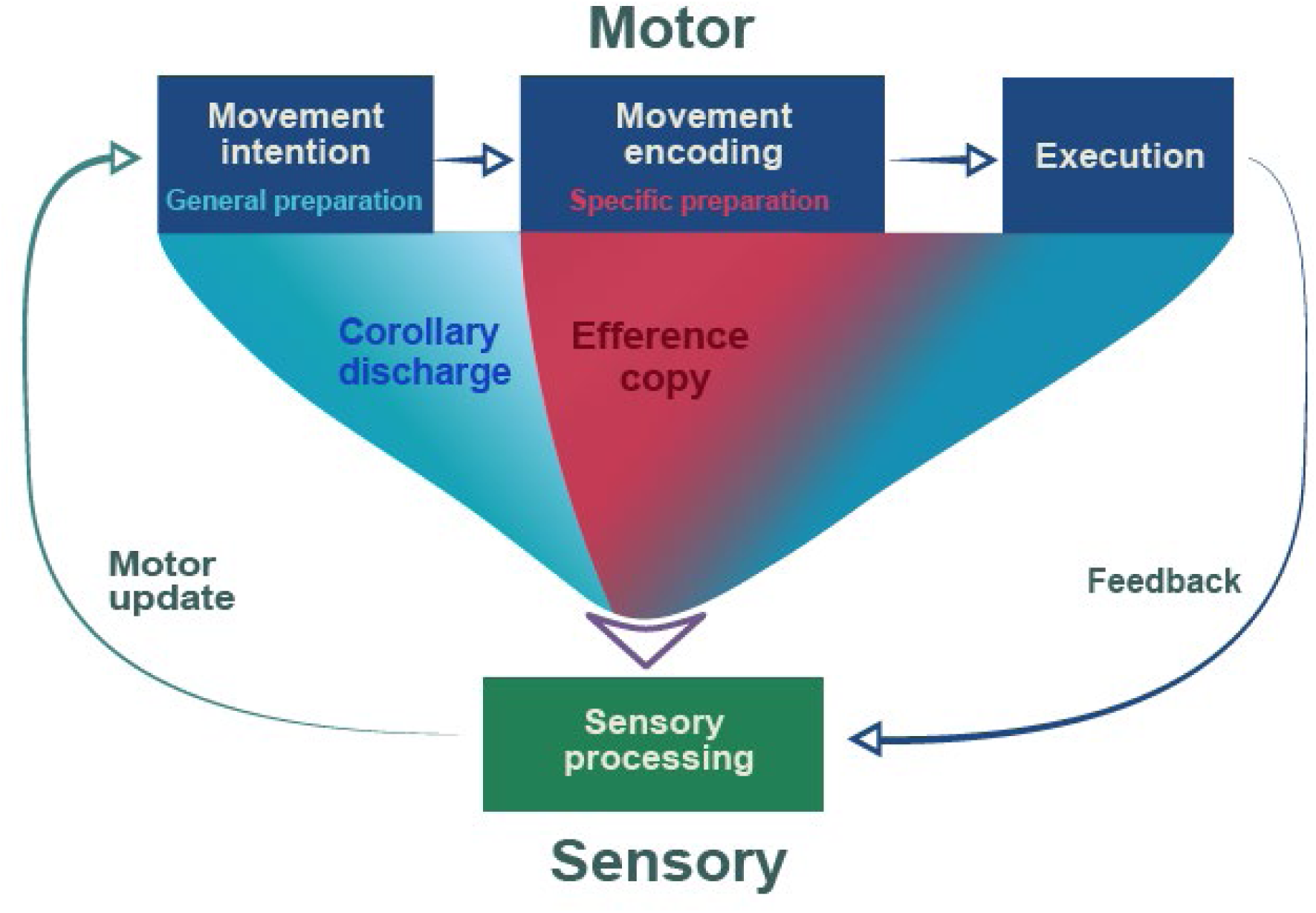
Schematics of proposed motor signals and their functions in the motor-to-sensory transformation. The intention and preparation stages before execution are mediated by the upper-stream motor circuitries that could potentially provide different motor signals. Copies of different motor signals are available to transmit to sensory regions at distinct temporal stages. The *corollary discharge* (*CD*) is a discharge signal within the established motor-to-sensory transformation pathways, and it would be available during the *general preparation* (*GP*) stage (in the movement intention phase). The *CD* does not necessarily include any content information. Its function could be inhibiting of all processes in the connected sensory regions, indicating the impending motor actions (cyan-shaded arrow). Whereas the *efference copy* (*EC*) would be available during the *specific preparation* (*SP*) stage (in the movement encoding phase) — after the development of a concrete movement plan. Its function could be selectively modulating the neural responses to the prepared syllable sounds (red-shaded arrow). The last execution stage could bundle all available motor signals and yield the observed mixed results and competing functions.

More importantly, we hypothesize that the functions of the distinct motor signals are determined by their contents. Similar to the arguments that a single type of corollary discharge would be too simplified to reflect the complexity of the motor signals regarding their sources, targets, and functional utilities (Crapse and Sommer 2008), we specify the putative functions by referring to the literal meanings of the two historical terms. Specifically, the *corollary discharge* is a discharge signal within the established motor-to-sensory transformation pathways. It does not necessarily include any content information. Its function could be, as manifested in saccadic suppression (Ross et al. 2001) and speech-induced suppression (Houde et al. 2002), inhibition of all processes in the connected sensory regions, indicating the impending motor actions (illustrated as a cyan-shaded arrow in Fig. 1). Whereas, the *efference copy* is an identical copy of motor signals that include detailed codes about actions. It is generated in a manner of one-to-one mapping in the motor-to-sensory transformation pathway. Its function could be, as indicated in the amplification of the mormyromast electroreceptors for electrolocation in electric fishes (Mohr et al. 2003) and priming the echo-sensitive neurons for echolocation in bats (Schuller 1979; Neuweiler 2003), selectively enhancing the sensitivity to reafferent (sensory feedback) caused by actions (illustrated as a red-shaded arrow in Fig. 1). Human speech data also offer hints suggesting the possible separation of these two functions, for example, the cortical sites of speech suppression and feedback sensitivity do not overlap (Chang et al. 2013). That is, we specify distinct functions in the otherwise interchangeably used historical terms to reflect our hypothesis that the functional specificity of motor signals is constrained by their contents. Together with the hypothesis about the distinct dynamics of these motor signals, the updated theoretical framework may account for the mixed neural modulations and competing functions of motor-to-sensory transformation.

In this study, we tested these hypotheses in the domain of human speech. The proposed functional specificity of the forward motor signals should be a canonical neural computation among animal species and across motor-related cognitive functions. However, because the nature of our hypotheses — the content information in different stages before action execution determines distinct functions of motor signals, the experimental manipulations on complex task requirements put a high demand on training animals. Therefore, we investigated these hypotheses with a novel delayed articulation paradigm using non-invasive human scalp electroencephalography (EEG) recordings. We targeted the early auditory responses of N1/P2 components to manifest the modulation effects of *CD* and *EC* on auditory processing. Previous studies have consistently demonstrated that speech production modulates early auditory responses (Behroozmand et al. 2009; Liu et al. 2011; Chen et al. 2012). Therefore, we hypothesized that the motor signals, which generate during speech preparation, would possess similar characteristics and influence early auditory responses.

In a series of three experiments, participants prepared to speak according to different visual cues. When the cues were symbols, participants generally prepared the action of speaking without any linguistic information (Experiments 1 & 2). According to our hypothesis, the *CD* would be available during this *general preparation* (*GP*) stage (as in the movement intention phase in Fig. 1) and would suppress neural responses to all sounds. When participants prepared to speak a syllable indicated by the written syllabic cues, the *EC* would be available during this *specific preparation* (*SP*) stage (as in the movement encoding phase in Fig. 1) and would selectively enhance the neural responses to the prepared syllable sounds (Experiment 1). The effects of the *CD* and *EC* would be independent of those of attention so that attentional effects on auditory processing (Experiment 3) would be different from the modulation effects of the motor signals (Experiments 1&2).

## Methods

### Participants

A total of 16 volunteers (5 males; mean age = 23.13; age range, 19-31 years) participated in Experiment 1; 19 participants (5 males; mean age = 23.89; age range, 19-31 years) in Experiment 2, and 17 participants (4 males; mean age = 23.94; age range, 20-35 years) in Experiment 3. All participants were right-handed native Mandarin speakers from East China Normal University. All participants had normal hearing without neurological deficits (self-reported). They received monetary incentives for their participation. Written informed consent was obtained from every participant. All protocols were approved by the institutional review board at New York University Shanghai, which was following the Declaration of Helsinki as a statement of ethical principles concerning human testing.

### Materials

Four audible syllables (/ba/, /ga/, /pa/, /ka/) and a 1k Hz pure tone with a duration of 400 ms were synthesized using the Neospeech web engine (www.neospeech.com) at a sampling rate of 44.1kHz in a male voice. All auditory stimuli were presented binaurally at 70 dB SPL, via plastic air tubes connected to foam earplugs (ER-3C Insert Earphones; Etymotic Research). A Shure beta 58A microphone was used to detect and record participants’ vocalization. Materials were consistent throughout three experiments.

### Procedure

#### Experiment 1: distinct preparation stages in a delayed-articulation task

Experiment 1 was an omnibus paradigm that included separate speech preparation stages before articulation. It aimed to prove the working principle of temporal segregating and inducing of *CD* and *EC* in different stages of action preparation. Auditory probes were introduced during each preparation stage to investigate the nature of the motor signals by testing how distinct preparation stages modulate the perceptual responses to the auditory probes.

We designed a delayed-articulation paradigm. Fig. 2A illustrates one type of trial in which participants were asked to produce a syllable according to visual cues after two stages of preparation. The trial started with a fixation cross displayed for 500 ms, followed by two consecutive preparation stages, each of which includes a visual cue that appeared in the center of the screen for a duration that jittered between 1500 ms to 2000 ms. Participants were instructed to make different preparations according to the cue. The visual cue in the first stage was meaningless symbols (#%) in yellow (blue in Fig.2*A* for better illustration) that did not contain any linguistic information (*general preparation*, *GP*). The visual cue in the second stage was a syllable in red that was randomly selected from /ba/, /ga/, /pa/, /ka/ (*specific preparation*, *SP*). The two preparation stages were separated by a blank with the duration jittered between 600 ms and 800 ms. During the last 400 ms of each preparation stage, either a 1k Hz pure tone or one of four auditory syllables (/ba/, /ga/, /pa/, /ka/) was presented to probe the modulatory function of preparatory motor signals. In *SP*, the auditory probe of syllables was either the same as or different from the visual cue, yielding two conditions -- auditory syllables were congruent with the visual information and hence the prepared syllable (*SPcon*) or incongruent (*SPinc*). After the offset of the sound in *SP*, a blank with the duration jittered between 600 ms and 800 ms was presented and was followed by a syllable in green that was the same as the written syllable in the *SP stage*. Participants were asked to produce the syllable as fast and accurately as possible. The onset times of vocal responses were recorded to quantify the reaction time.

**Figure 2.**
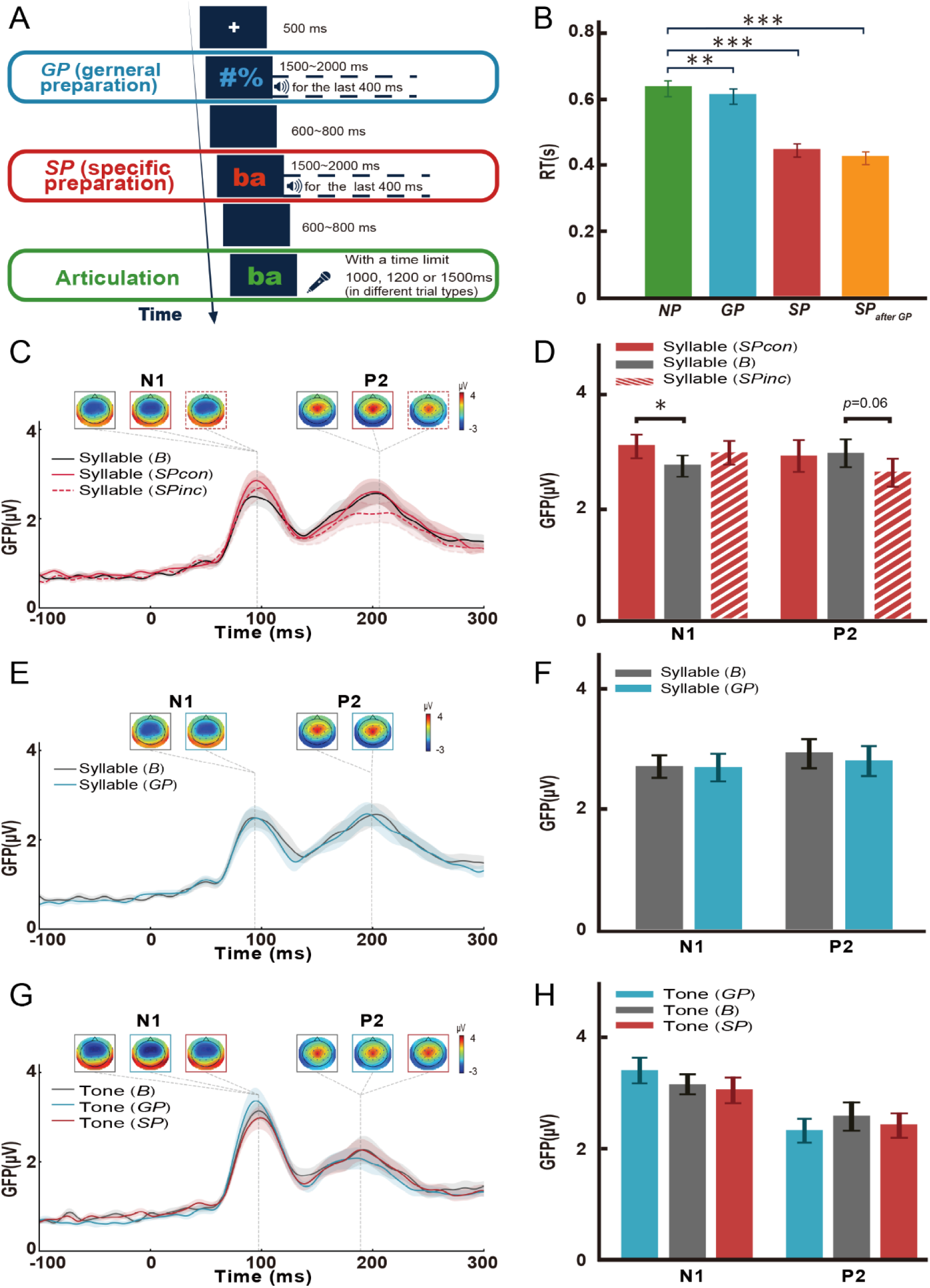
Experimental paradigm, behavioral, and ERP results of Experiment 1. **A)** Illustration of a sample trial that includes all preparation stages. Participants were asked to prepare to articulate a syllable according to visual cues that were either symbols (*general preparation, GP –* preparing to speak without knowing the content) or syllables in red (*specific preparation, SP –* preparing to speak the specific content). When a syllable in green appeared, participants were required to pronounce it rapidly. An auditory probe (a 1k Hz pure tone or a syllable sound) was presented during each preparation stage. Additional types of trials were included by randomly combining the preparation stages and the articulation tasks to provide better control of the preparation and to yield baseline responses. (Refer to Methods for all types of trials and conditions.) **B)** The speed of pronunciation was measured as reaction time (RT). Error bars indicate +- SEM. \*\**p* < 0.01, \*\*\**p* < 0.001. Faster articulation speed on *GP* and *SP* conditions than the *NP* condition. **C)** ERP time course and topographic responses for *SP* and *B* conditions. Individual peak amplitudes and peak latencies for the N1 and P2 GFP waveform were observed in each condition. The response topographies at each peak time are shown in colored boxes near each peak, using the same color-coding to represent each condition. The *SP* enhanced the N1 responses to the prepared syllables (*SPcon*). **D)** Mean GFP amplitudes across participants at N1 and P2 latencies for *SP* (red bars) and *B* (grey bars) conditions, respectively. *SP* enhanced the N1 responses to the prepared syllables (*SPcon*). Error bars indicate ± SEMs. Asterisks show the significance of post hoc *t-tests*, FDR-corrected for multiple comparisons (\**p* < 0.05). **E)** ERP time course and topographic responses for *GP* and *B* conditions show that no modulation effects of the *GP* on N1 and P2 responses to syllables. **F**) Mean GFP amplitudes across participants at the N1 and P2 latencies for *GP* (blue bars) and *B* (grey bars) conditions as observed in **E**. **G**) No modulation effects of *GP* and *SP* on the N1 and P2 responses to tones. **H**) Each bar represents the mean GFP amplitudes across participants at N1 and P2 latencies for each condition to tones. The red bars depict the *SP* condition, the blue bars depict the *GP* condition, and grey bars depict the *B* condition.

To better control preparation and reduce expectation, in addition to the type of trials that included *GP*, *SP*, and articulation in a sequence as described above (*SPafter GP trials*, Fig. 2A), visual cues were pseudorandomly paired and yielded three more types of trials. First, the green syllable articulation cue could immediately appear after the fixation (immediate vocalization without preparation, *NP trials*). The reaction time in *NP* trials served as a baseline behavioral response of syllable production and compared with reaction times in other trials to quantify the effects of preparation behaviorally. Second, the articulation cue could follow the general preparation cue (*GP trials*). Third, the specific preparation cue could appear directly without GP (*SP trials*). Therefore, a total of four types of articulation task trials were included. The time limits for articulation were set to 1500 ms, 1200 ms, 1000 ms, and 1000 ms in *NP*, *GP*, *SP*, and *SP*_*after GP*_ trials, respectively. These manipulations were to eliminate any expectations and enforce preparation.

Moreover, in another type of trial, participants saw a white visual cue (**) without any linguistic information. No articulation green syllable cues followed the white symbols in these trials. Participants only needed to passively listen to the auditory probes without the requirement of action preparation or articulation (baseline listening without preparation, *B*). The *B* trials, which had similar visual cues and auditory probes but without preparation, yielded baseline auditory responses to quantify the neural modulation effects of preparation. Therefore, five types of trials (*NP, GP*, *SP*, *SP*_*after GP*_, & *B*) were randomly presented in five blocks. In each block 64 trials were included, yielding a total of 320 trials in the experiment, with 60 trials for each type of the *NP, GP*, *SP*, and *SP*_*after GP*_ trials and 80 trials for *B*. The number of auditory probes in the *GP* and *SP* stages was 120 each, and each of the *SPcon* and *SPinc* conditions had 60 auditory probes.

#### Experiment 2: probabilistic auditory probes enforcing the general preparation

We varied the duration of visual cues to eliminate the temporal expectation of auditory cue onset time in Exp. 1. However, the auditory probes were always following the visual cue. This temporal association could grant participants a strategy that they could start to prepare after hearing the auditory probe. That is, the motor signals of interest were not induced throughout the preparation stages, which seriously dampened the modulation effects to the auditory probes, especially in the *GP* conditions, as the null results in Exp.1 indicated. Experiment 2 aimed to control this confound by introducing trials that did not contain auditory probes. The mixed trials enforced participants to prepare to speak according to the visual cues even though they did not know what syllable to speak. This experimental manipulation increased the power to investigate the functions of *CD* during *GP*.

The experiment procedure was similar to the one in Experiment 1, except that only the general preparation task (*GP*) was included. Half of the trials did not include the auditory probe (*GPNS*), as illustrated in Fig. 3*A*. Such mixed trials enforced participants to prepare the final vocalization task based on the visual cues instead of auditory stimuli. The time limit for the articulation task was set to be 1500 ms for *NP*, 1200 ms for *GP* and *GP*_*NS*_ trials, respectively. Four types of trials (*NP, GP*, *GPNS*, *B*) were randomly presented in four blocks. In each block, 64 trials were included, yielding a total of 384 trials in the experiment, with 96 trials in each of the *NP, GP*, *GP*_*NS*_, and *B*. Each of the *GP* and *B* conditions had 96 auditory probes.

**Figure 3.**
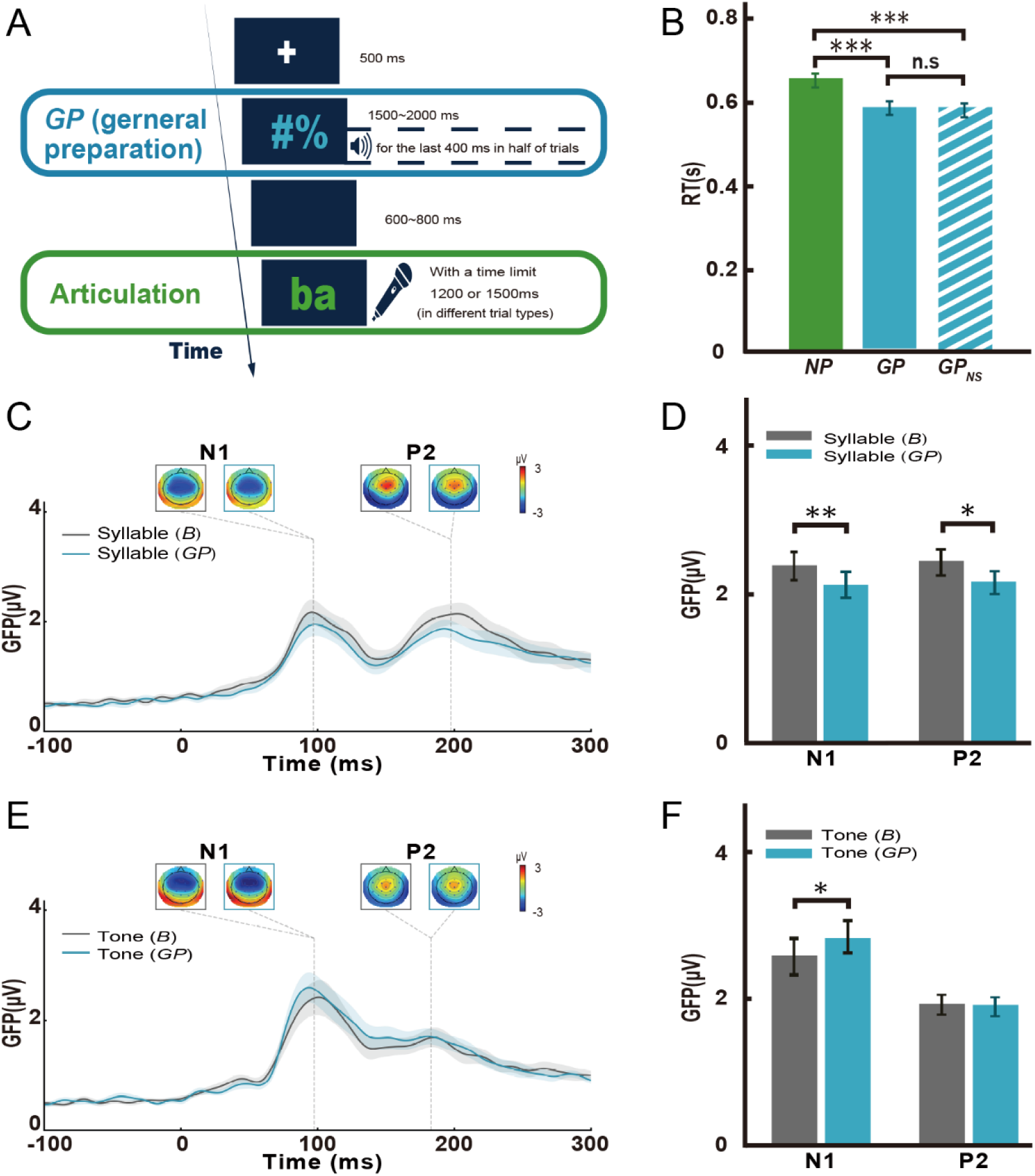
Experimental paradigm, behavioral, and ERP results for Experiment 2. **A)** Experiment 2 is similar to Experiment 1 except that participants performed the *GP* condition only. Half trials were without the auditory probes so that the mixed trials enforced participants to prepare to speak based on the visual cues without knowing the speech content. **B)** Facilitation in reaction time by the general preparation with (*GP*) or without sound probes (*GP*_*NS*_), *** *p* < 0.001, *n.s*: not significant. **C)** ERP waveforms and topographic responses for *GP* and *B* conditions. The response topographies at each peak time are shown in colored boxes near each peak, using the same color-coding to represent each condition. Suppression in N1 and P2 responses to syllable sounds by the *GP* was observed. **D)** Mean GFP amplitude across participants at N1 and P2 latencies for *GP* (blue bars) and *B* (grey bars) conditions as observed in **C**, ** *p* < 0.01, * *p* < 0.05. **E)** Enhancement in N1 responses to tones by the *GP.* **F)** Each bar represents the mean GFP amplitudes across participants at N1 and P2 latencies for each condition to tones, respectively. The blue bar depicts the *GP* condition and grey bar for the *B* condition, * *p* < 0.05.

#### Experiment 3: explicitly directing attention to auditory probes during preparation

Arguably, during preparation, attention is shifted to the perceptual consequences of actions. In Experiments 1 & 2, when preparing to speak, participants were likely to direct their attention to the sound that they were going to produce. Therefore, the observed modulation effects on auditory probes could be induced by attention. However, because it is hard, if not impossible, to completely wipe out attention, we explicitly direct participants’ attention to the auditory probes by a task related to the auditory probes in this experiment. If the observations in Experiments 1 & 2 were caused by the attention, we should obtain similar results in this experiment. Otherwise, the results in previous experiments cannot be accounted for by attention.

The experiment procedure is similar to Experiment 1, except that participants were required to identify the auditory probes. During the general preparation task, participants were asked to identify the upcoming auditory probe, whether it was a syllable or a tone. During the specific preparation task, participants were asked to determine whether the visual cue and auditory probe were congruent or incongruent. Participants needed to make the identification response within a time limit of 2000 ms (Fig. 4*A*). Five types of trials (*NP, GP*, *SP*, *SP*_*after GP*_, & *B*) were randomly presented in five blocks. In each block 64 trials were included, yielding a total of 320 trials in the experiment, with 60 trials for each type of the *NP, GP*, *SP*, and *SP*_*after GP*_ trials and 80 trials for *B*. The number of auditory probes in the *GP* and *SP* stages was 120 each, and each of the *SPcon* and *SPinc* conditions had 60 auditory probes.

**Figure 4.**
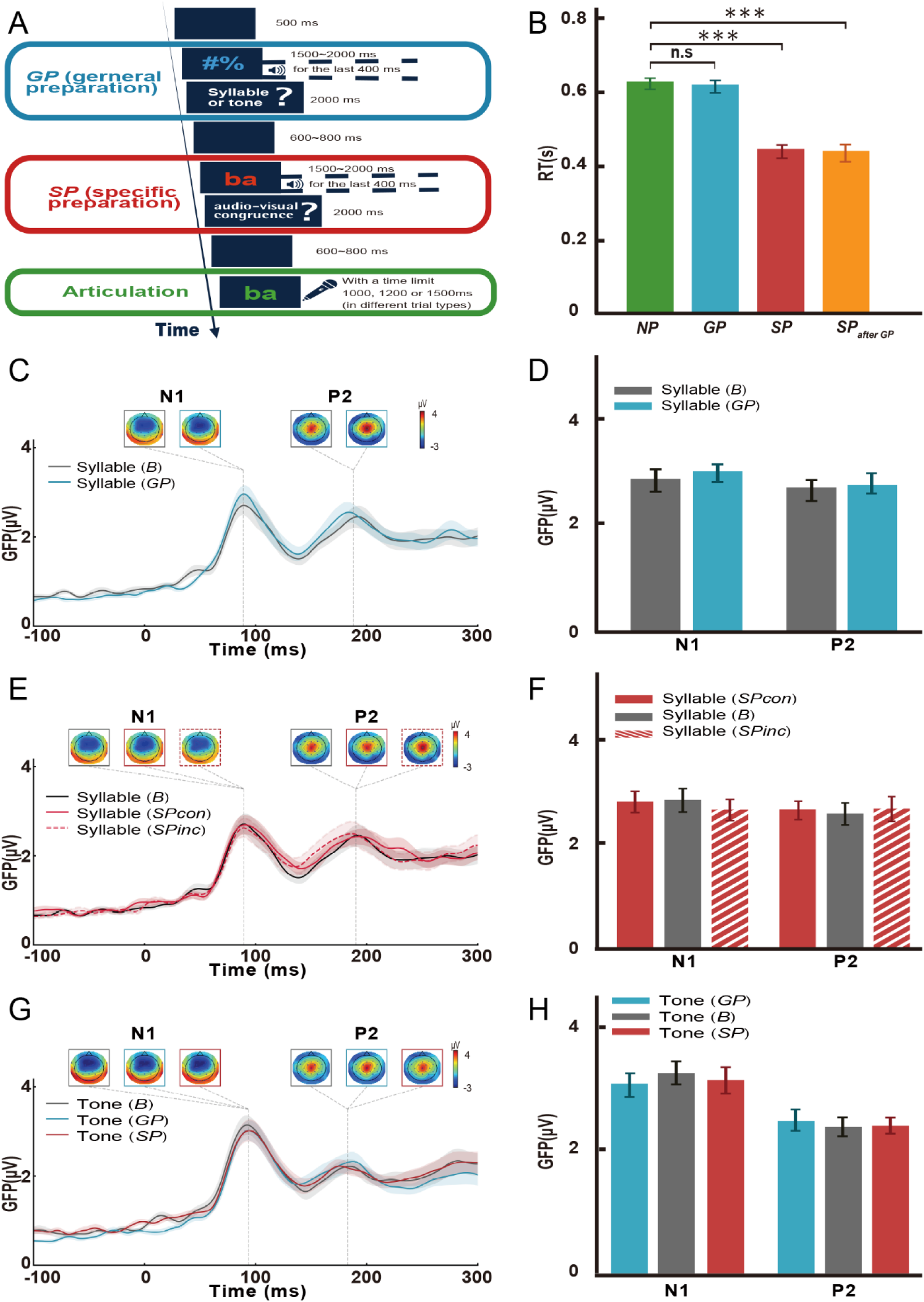
Experimental paradigm, behavioral, and ERP results for Experiment 3. **A)** Participants were explicitly instructed to identify the auditory syllables. During *GP*, participants were asked to identify the upcoming auditory probe, whether it was a syllable or a tone. During *SP*, participants were asked to determine whether the visual cue and auditory probe were congruent. **B)** Facilitation in reaction time by *SP*. \*\*\**p* < 0.001. **C)** ERP time course and topographic responses for *GP* and *B* conditions. Individual peak amplitudes and peak latencies for the N1 and P2 GFP waveform were observed in each condition. The response topographies at each peak time are shown in colored boxes near each peak, using the same color-coding to represent each condition. The N1 and P2 responses for syllables in the *GP* condition are not significant. ***D***, Mean GFP amplitudes across participants at N1 and P2 latencies for *GP* (blue bars) and *B* (grey bars) conditions as observed in **C**. **E)** The effect was not significant in both the N1 and P2 response in the *SP* condition for syllables. **F**) Each bar represents the mean GFP amplitudes across participants at the N1 and P2 latencies for each condition to tones, respectively. The red bars depict the *SP* condition and grey bar the *B* condition. **G**) No effects on the N1 and P2 responses to tones in both *GP* and *SP* conditions. **H**) Each bar represents the mean GFP amplitudes across participants at N1 and P2 latencies for each condition to tones. The red bars depict the *SP* condition, the blue bars depict the *GP* condition, and grey bars depict the *B* condition.

### Data analysis

#### Behavioral data analysis

The reaction times (RTs) of the articulation task were calculated as the time lag between the onset of the green visual cue and the onset of the vocalization. In Experiment 1, averaged RTs were obtained in each of the four trial types (*GP*, *SP*, *SP*_*after GP*_, *NP*). The RT data was subject to a repeated-measures one-way ANOVA and a post-hoc Tukey Student *t-test* for pairwise comparisons. Behavioral data analysis in Experiment 2 was similar to Experiment 1 except that the reaction times from trials were averaged in each of the three trial types (*NP*, *GP*, *GP*_*NS*_). The same statistical methods were applied. In Experiment 3, behavioral data analysis was identical to Experiment 1, averaged RTs were obtained in each of the four trial types (*GP*, *SP*, *SP*_*after GP*_, *NP*).

#### EEG data acquisition and processing

Neural responses were recorded using a 32-channel active electrode system (Brain Vision actiCHamp; Brain Products) with a 1000 Hz sampling rate in an electromagnetically shielded and sound-proof room. Electrodes were placed on an EasyCap, on which electrode holders were arranged according to the 10-20 international electrode system. The impedance of each electrode was kept below 10 kΩ. The data were referenced online to the electrode of Cz and re-referenced offline to the average of all electrodes. Two additional EOG electrodes (HEOG and VEOG) were attached for monitoring ocular activity. The EEG data were acquired with Brain Vision PyCoder software (http://www.brainvision.com/pycorder.html) and filtered online between DC and 200 Hz with a notch filter at 50 Hz.

EEG data processing and analysis were conducted with customized Python codes, MNE-python (Gramfort et al. 2014), EasyEEG (Yang et al. 2018), and TTT toolboxes (Wang et al. 2019). For each participant’s dataset, noisy channels were manually rejected during visual inspection. The dataset was band-pass filtered with cut-off frequencies set to 0.1 and 30 Hz. The filtered dataset was then cut into epochs ranging from −200 ms to 800 ms, relative to the onset of the auditory probe, and baseline corrected using the 200 ms pre-stimulus period. Epochs with artifacts related to eye blinks and head movement were manually rejected. Epochs with peak-to-peak amplitude exceeded 100 *µ*V were automatically excluded. To ensure data quality, we excluded epochs before analysis if they were contaminated by any residual noise. The remaining epochs were used to obtain the average event-related responses (ERP) in each condition. An average of 244 (*SD* = 42.3) epochs for each participant were included in Experiment 1, 208 (*SD* = 22.0) epochs in Experiment 2, and 233 (*SD* = 28.7) epochs in Experiment 3. The number of trials in each condition of Experiments 1, 2, and 3 is as follows. In Experiment 1, on average, 31, 43, 31, and 31 trials were included for conditions with auditory syllables in *B*, *GP*, *SPcon*, *SPinc*, respectively. For conditions with tones, on average, 30, 45, and 31 trials were included in *B*, *GP*, *SP*, respectively. In Experiment 2, on average, 36 and 35 remaining trials were in *B* and *GP*, respectively (same number for both syllables & tones). In Experiment 3, for auditory syllables, on average, 30, 43, 28, and 29 trials were included in *B*, *GP*, *SPcon*, *SPinc*, respectively. For conditions with tones, on average, 40, 35, and 29 trials were included in *B*, *GP*, *SP*, respectively. The ratio of trial-rejection in Experiments 1, 2, and 3 was 23.75%, 26.04%, and 27.18%, respectively. To conclude, we rejected ~25% of trials for each experiment.

In Experiment 1, the global field power (GFP) -- the geometric mean across 32 electrodes -- was calculated separately for tones in three conditions (*GP*, *SP*, and *B*), and for the auditory probes of syllables in four conditions (*GP*, *SPcon*, *SPinc*, and *B*). An omnibus measure such as GFP is optimal in a novel study to balance between the requirements of exploration and problems of false positives by avoiding subjective channel selections, multiple comparisons, and individual differences (Tian and Huber 2008; Tian et al. 2011; Yang et al. 2018; Wang et al. 2019). Individual peak amplitudes and peak latencies for the N1 and P2 components in the GFP waveforms were automatically identified using the TTT toolbox in predetermined time windows of 90 to 110 ms and 190 to 210 ms, respectively (Wang et al. 2019). We visually verified that identified peaks by the toolbox were within the correct time windows in each participant. For responses to syllables, paired *t*-tests were carried out between the comparisons of *GP* and *B*, *SPcon* and *B*, and *SPinc* and *B*, separately for the N1 and P2 components. For responses to tones, repeated-measures one-way ANOVAs were conducted among the responses to the auditory probes in *GP*, *SP*, and *B*, separately for the N1 and P2 components. For ANOVAs, effect sizes were indexed by partial *η*^*2*^. For paired t-tests, effect sizes were indexed by Cohen’s d. We calculated these effect sizes indexes to quantify the proportion of variance. Significant effects were determined by *p* < 0.05 and partial *η*^*2*^ >0.14 (Richardson 2011).

In Experiment 2, EEG data analysis was similar to Experiment 1. For responses to syllables, paired *t*-tests were carried out between *GP* and *B* conditions, separately for the N1 and P2 components. For the *GP*_*NS*_ condition, epochs were extracted around a similar latency as the *GP* condition. The same peak selection and ERP analysis methods were used to determine the exact peak latency to verify that no auditory responses were induced in *GP*_*NS*_. Statistical methods were similar to Experiment 1. Both for responses to syllables and tones, paired *t*-tests were carried out between *GP* and *B* conditions, separately for the N1 and P2 components.

In Experiment 3, EEG data processing was identical to Experiment 1. For responses to syllables, paired *t*-tests were carried out between *GP* and *B* conditions, as well as comparing *SPcon* and *SPinc* conditions to *B*, separately for the N1 and P2 components.

For responses to tones, repeated-measures one-way ANOVAs were conducted among the responses to the auditory probes in *GP*, *SP*, and *B*, separately for the N1 and P2 components.

### Modeling

To quantify the proposed distinctions between *CD* and *EC*, we built a two-layer neural network model to simulate the dynamics and modulation effects of motor signals on sensory processing (Fig. 5A). The model structure and basic units are similar to previous models (Huber and O’Reilly 2003; Ma and Tian 2019). The upper layer represents motor processing, and the lower layer represents auditory processing. Each layer includes multiple neurons that represent different syllables. (Only four nodes are drawn for illustration purposes.) Each neuron in the auditory layer is a rate-coded unit with synaptic depression. The updating of membrane potential is governed by Eq. 1.

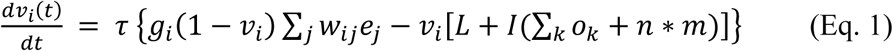

**Figure 5.**
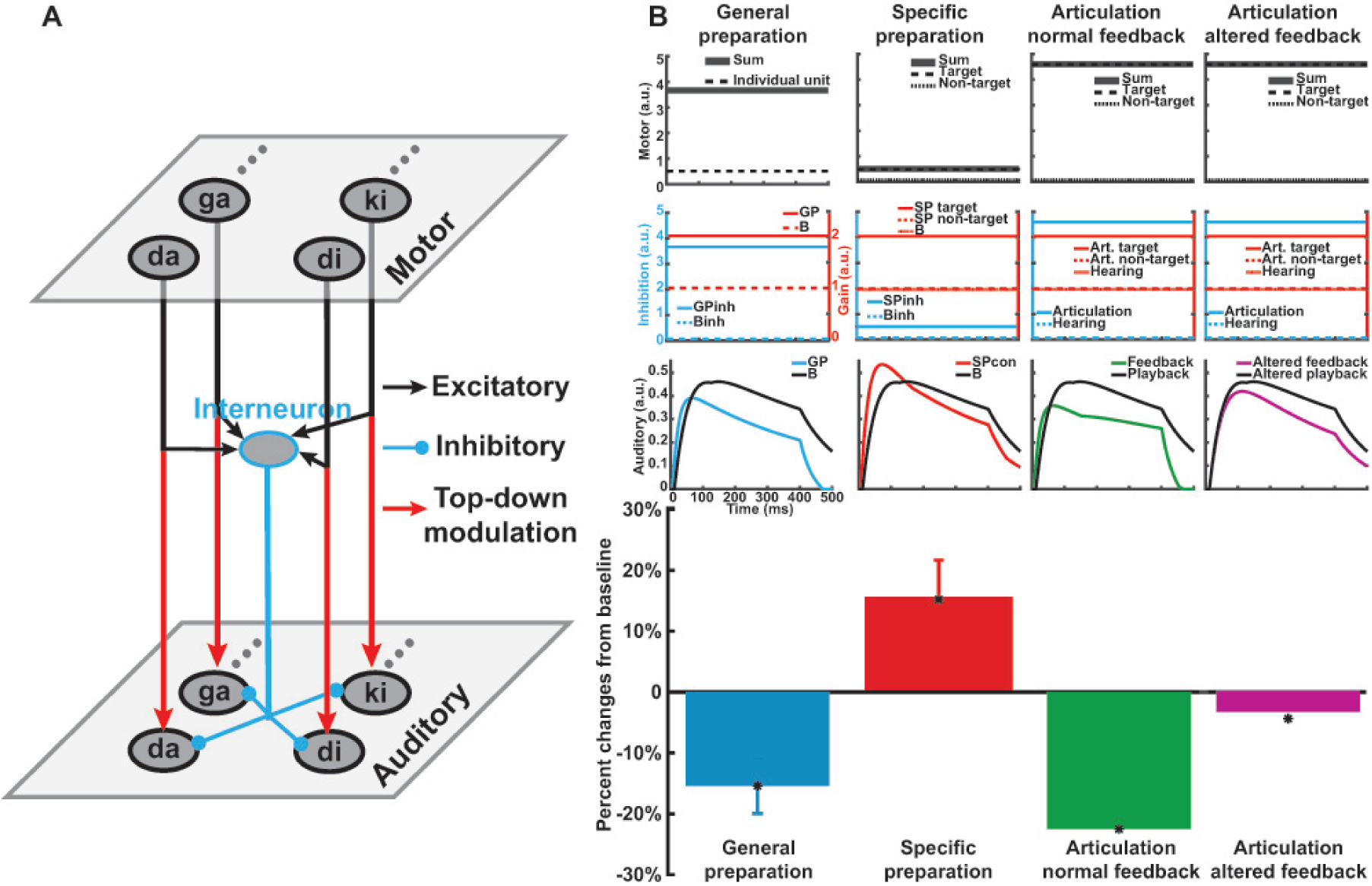
A neural network model of distinct motor signals and simulation results. **A)** Bifurcation of motor signals realizes distinct functions in a neural network model. A motor layer and an auditory layer include nodes that represent syllables. Each node is a rate-coded leaky-integrate-and-fire neuron. Signals from each motor unit split into two. One branch of the signals directly modulates the post-synaptic gain of the corresponding auditory unit, simulating the function of *EC* (the red line). The other branch of the signals accumulates and actives an interneuron that inhibits all auditory units, simulating the function of *CD*. **B)** Simulation results capture the modulation dynamics in speech preparation and execution. Four simulations of modulation effects in different stages during speech production are arranged in columns, including general preparation (the first column), specific preparation (the second column), articulation with normal feedback (the third column), and articulation with altered feedback (the fourth column). Plots in the first row represent the motor activity in an individual motor unit as well as the sum from all units. Plots in the second row show the inhibitory strength and modulation gain on the auditory units. Plots in the third row depict the dynamics of auditory responses in different conditions. The bottom row includes empirical and simulation results. Bars are empirical data after converting into percent changes [(experimental condition - baseline)/baseline]. The first two bars are modulation results of the general and specific preparations in Fig. 3D and Fig. 2D, respectively. The last two bars are speech-induced suppression and reduction of suppression to altered feedback by averaging the effects from the left and right hemispheres (Houde et al. 2002). Stars represent the simulation results. In the first column, the lack of detailed information during the general preparation stage causes activation in all motor units and results in suppression of all auditory units, yielding the observed auditory suppression in GP. In the second column, the detailed information available during the specific preparation stage activates a given motor unit and increases the sensitivity to the corresponding auditory unit, which yields an enhancement effect. In the third column, stronger inhibitory signals from a given motor unit during the execution of speech strongly inhibit the corresponding auditory unit, resulting in the commonly observed speech-induced suppression. In the fourth column, altered speech feedback input into non-target auditory units. Despite the same articulation (the same parameters as in the third column), the auditory suppression is reduced.

The member potential of neuron *i* at the auditory layer, *v*_*i*_, is updated according to the integration rate (time constant, *τ*) summing over three sources of input. The first input is an excitatory input from acoustic signals, *e*_*j*_, via bottom-up connection strength *w*_*ij*_. This bottom-up input drives the membrane potential to 1 (governed by the multiplier of 1-*v*). The second input is the leak with the fixed term *L*. The third input is the inhibition, which is the strength of *I* multiplied by the sum of two terms. One is the lateral inhibition that is the sum of output at time *t* from *k* units at the auditory layer. Another is the inhibition from the motor layer, *n*m*, which is specified next. The combination of the leak and inhibition drives the membrane potential towards 0 (as the term in the bracket is multiplied by -*v*). The fixed parameters are similar to those used in previous studies (Huber and O’Reilly 2003; Ma and Tian 2019).

The influences of motor signals are modeled as two sets of free parameters. The motor signals come from the same motor units but split into two sources. One source includes activities of all motor units integrated into an interneuron that inhibits all neurons in the auditory layer. For simplification, the inhibition from each neuron in the motor layer is assigned as a unit value, *m*. The equivalent inhibition effects from the interneuron are the sum of *n* motor units, *n*m*. This motor source simulates the hypothesized function of *CD*. Another source is the direct modulation between the corresponding syllable in two layers. This motor signal is modeled as a gain control parameter, *gi*, which increases the gain of the corresponding auditory unit to the excitatory input. This motor source simulates the hypothesized function of *EC*.

The implementation of an interneuron and gain modulation is motivated by previous research. Speech production and control models assume that internal forward models generate predictions that cancel the auditory processes of normal speech feedback (Guenther 2006; Hickok et al. 2011; Houde and Nagarajan 2011; Hickok 2012). Recent studies found that interneurons mediate these inhibitory functions (Schneider et al. 2014; Attinger et al. 2017). Moreover, previous studies suggest that predictions can be more precise than attention and provides more considerable gain for neural processing (Kok et al. 2012). Our empirical and modeling works have also suggested that the motor-based prediction via internal forward models in speech production are more precise than memory retrieval (Tian and Poeppel 2013; Tian et al. 2016) and provides a larger excitatory gain for predictive auditory feedback (Ma and Tian 2019). A Bayesian framework also suggests that updating prior, functionally similar to the gain modulation in neural network models, as the modulation effects of prediction (Aitchison and Lengyel 2017). Therefore, in this study, we combined the implementation of an interneuron and gain modulation in one neural network model to collaboratively simulate the inhibition and enhancement throughout the time course of speech production.

During the simulation of specific preparation (*SP*), only the prepared syllable in the motor layer is activated. Only one unit input, *m*, from the motor layer is sent to the interneuron. Moreover, a specific gain modulation is applied to the auditory neuron of the prepared syllable, *g*_*i*_. We first fitted the model to the observed enhancement during *SP* by adjusting the parameters of *m* and *g*_*i*_. Next, during the simulation of general preparation (*GP*), because of the lack of linguistic information, the preparation induces weak activates in all motor neurons, *n*. The sum of all motor units *n*m* would provide a more potent activity to the interneuron for inhibiting auditory processes. We fitted the model to the observed suppression in *GP* by adjusting the parameter of *n*.

The primary reason for constructing this model is to explain the observed preparation modulation results. To connect to the previous articulation suppression data, we assume that additional motor neurons in the downstream of speaking action beyond preparation are active. The motor activation can bring a strong inhibition as the auditory suppression occurs near the onset of speech production (Daliri and Max 2016). With the additional motor activity, the inhibition to the speech feedback could become much stronger and induce strong suppression to normal feedback compared to passive listening of playback. We fitted our model to the speech-induced suppression data in Houde et al. (Houde et al. 2002) by adjusting the additional motor activity. Last, to further test the model after fixing all free parameters, we examined whether the model can generate similar responses to the altered feedback as the empirical observations. We fed the acoustic input to the auditory unit that was next to the unit of speech target and a unit representing tones while having the identical procedure and parameters as in the last simulation of speech-induced suppression. We tested whether our model can generate a similarly reduced suppression, as observed in Houde et al. (Houde et al. 2002), when the tones were mixed in the normal feedback.

To assess how the model fitted the empirical results, we treated the model stimulation results as the mean from a distribution with unknown variance. Data in *GP* and *SP* conditions were subject to one-sample *t*-tests against the simulation results. Because this analysis was to test the null hypothesis that the model simulation results were from a similar distribution of empirical results, we used a Bayesian analysis method for one-sample *t*-tests (Rouder et al. 2009). (online tool at http://pcl.missouri.edu/bf-one-sample). The Bayes factor B01 = M0/M1, where M0 and M1 are the marginal likelihood for the null and alternative, respectively. That is, the Bayes factors are odds ratios between the null and alternative hypotheses, which means that the null is B01 times more probable than the alternative. The parameters we input for the Bayesian analysis were a sample size of 19 and 16 for *GP* and *SP* conditions and scale r on an effect size of 0.707. For testing the model fitting in the speech-induced suppression and reduction of suppression for altered feedback, because we do not have access to the raw data, we compare the simulation results to the 95% confidence intervals in the figures of Houde et al. (Houde et al. 2002).

## Results

### Experiment 1: distinct preparation stages in a delayed-articulation task

In Experiment 1, participants were asked to speak a syllable after various stages of preparation. A repeated-measure one-way ANOVA on RTs found a significant main effect of preparation (*F*(3,45) = 97.720, *p*<0.0001, partial *η*^*2*^ = 0.867). Further paired *t*-tests revealed that the onset of articulation was consistently faster after preparation. Specifically, articulation after *GP* (mean RT of 609 ms) was faster than immediate vocalization without preparation (*NP*, mean RT of 633 ms), (*t*(15) = 3.177, *p* < 0.01, *d* = 0.267). RTs were much shorter after *SP* (445 ms) than *NP* (*t*(15) = 10.678, *p* < 0.0001, *d* = 2.164). RTs were shortest when they articulated after *GP* and *SP* in a row (*SP*_*after GP*_: 422 ms) (*t*(15) = 10.584, *p* < 0.0001, *d* = 2.512) (Fig. 2B). The *GP* before *SP* did not provide additional facilitation, as the RT difference between *SP* and *SP*_*after GP*_ was not significant (*t*(15) = 0.607, *p* > 0.05, *d* = 0.053). These behavioral results suggested that participants engaged in speech preparation.

We further scrutinized the EEG neural responses to investigate the functions of motor signals during preparation. Paired *t*-tests were carried out between the auditory responses to the probes in the general preparation (*GP*) and specific preparation (*SP*) conditions, separately for the N1 and P2 components. In the *SP*, early neural responses of N1 were larger than that in *B* when the auditory syllables were congruent with the specific preparation visual cues (*SPcon*) (*t*(15) = −2.49, *p =* 0.025, *d* = −0.432). However, the effect was not significant in the later auditory responses of P2 (*t*(15) = 0.248, *p* = 0.808, *d* = 0.039). The effects for the auditory syllables that were incongruent with the visual cue (*SPinc*) showed an opposite pattern. The effect in N1 was not significant (*t*(15) = −1.48, *p* = 0.160, *d* = −0.283), nor in P2 (*t*(15) = 2.024, *p* = 0.061, *d* = 0.342) (Fig. 2*D*). These results suggested that motor signals during specific preparation modulated the perceptual responses based on the content congruency.

In the *GP*, responses to auditory syllables were not different from the ones without preparation (*B*) neither in N1 (*t*(15) = 0.077, *p* = 0.940, *d* = 0.017), nor in P2 (*t*(15) = 1.070, *p* = 0.301, *d* = 0.119) (Fig. 2*F*). These results contrast with the ones obtained in the *SP*, presumably because motor signals with different natures were induced during distinct preparation stages. However, the null results in *GP* were different from what we predicted -- the *corollary discharge* that was induced during *GP* would suppress auditory responses. We speculated that the auditory probes, which were always presented at the last period of the preparation stage, would be a potential problem in this omnibus paradigm. That is, the general preparation could start toward the end of the stage so that the modulatory power of *corollary discharge* was significantly dampened. We addressed this potential problem in Experiment 2.

For responses to tones, repeated-measures one-way ANOVAs were conducted among three conditions (*GP*, *SP*, and *B*) for N1 and P2 separately. The effect was not significant neither in the early auditory responses of N1 (*F*(2,30) = 2.894, *p* = 0.07, partial *η*^*2*^ = 0.162) nor in the later auditory response of P2 (*F*(2,30) = 2.111, *p* = 0.138, partial *η*^*2*^ = 0.123). These results showed that no modulation effects of motor signals on tones during either preparation stages (Fig. 2*H*). These results of tones contrasted with the results of auditory syllables, indicating the motor signals during preparation contained the task-related information. In summary, the results of Experiment 1 suggested that different motor signals were generated during distinct preparation stages and modulated perceptual neural responses based on the contents of signals.

### Experiment 2: probabilistic auditory probes enforcing general preparation

The temporal association between the visual cues and auditory probes in *GP* could dampen the effects of *corollary discharge* and cause the null results in Experiment 1. In this experiment, we added trials without auditory probes during *GP* so that participants must prepare to speak according to the visual cues without linguistic information. The behavioral data showed a significant main effect of preparation (*F*(2,36) = 105.101, *p* < 0.0001, partial *η*^*2*^ = 0.854). Further paired *t*-tests revealed that RTs were facilitated when participants performed *GP* with sound probe than immediate articulation *NP* (mean RT for *NP* 651 ms; *GP*, 580 ms; *t*(18) = 10.534, *p* < 0.0001, *d* = 0.971). These results replicated the observations obtained in Experiment 1. More importantly, RTs were also significantly shorter when participants performed *GP* without sound probes than immediate articulation *NP* (mean RT for *GP*_*NS*_ 586 ms; *t*(18) = 11.078, *p* < 0.0001, *d* = 0.893). These results suggested that participants performed the *GP* task according to the visual cues and ensured that *corollary discharge* was available throughout the *general preparation* stage (Fig. 3*B*).

For ERP responses to the auditory probes of syllables, paired *t-*tests revealed that the amplitude of early N1 response in *GP* was less than that in *B* (*t*(18) = 3.406, *p* = 0.003, *d* = 0.3). The amplitude of later P2 response in *GP* was reduced relative to *B* (*t*(18) = 2.240, *p* = 0.038, *d* = 0.342) (Fig. 3*D*). The topographies were consistent in both conditions (Fig. 3*C*). These results, obtained after presenting the auditory probes in a probabilistic manner and increasing the power of *corollary discharge*, were consistent with our hypothesis -- *corollary discharge* that was induced during general preparation suppressed auditory responses.

Paired *t*-tests were also carried out on the auditory responses to the tones. Significantly larger N1 amplitude was revealed during *GP* compared with that in *B* (*t*(18) = −2.397, *p* = 0.028, *d* = −0.217). The effect was not significant in the later auditory response of P2 (*t*(18) = 0.194, *p* = 0.849, *d* = 0.028) (Fig. 3*F*). The enhancement in *GP* to the tones contrasted with the suppression results for the syllables, suggesting that the *corollary discharge* was motor-specific to actions -- speech production in this experiment. Violation of the goal of actions (e.g., pure tones that were not adapted to human vocal tracks and articulators), would create an error term and reverse the suppression effects. In summary, the results in Experiment 2 indicated the suppressive function of *corollary discharge*, which may be constrained by the task demand.

### Experiment 3: explicitly directing attention to auditory probes during preparation

The modulation effects observed in Experiments 1 & 2 could be due to the shift of attention to the prepared speech sounds. However, it is hard, if not impossible, to disentangle motor preparation and attention. In this experiment, we explicitly instructed participants to identify the auditory probes during the *GP* and *SP* to examine whether the attentional effects differ from previous observations in Experiments 1 & 2. All participants accomplished the identification task (accuracy of every participant was above 90%).

A repeated-measure one-way ANOVA on the articulation RTs revealed a significant main effect of preparation (*F*(3,48) = 51.020, *p* < 0.0001, partial *η*^*2*^ = 0.76). A further paired *t*-test revealed that RTs were shorter in *SP* than immediate articulation *NP* (*NP*: 621 ms, *SP*: 441 ms, *t*(16) = 8.315, *p* < 0.0001, *d* = 2.593). RTs were shortest after having both *GP* and *SP* (*SP*_*after GP*_: 437 ms) than immediate articulation *NP* (*t*(16) = 7.234, *p* < 0.0001, *d* = 2.318). However, the RT difference between *GP* and *NP* was not significant (*GP*: 614 ms, *t*(16) = 1.680, *p* = 0.112, *d* =0.110) (Fig. 4*B*). Overall, the behavioral results were consistent with the findings in Experiment 1.

The effects in neural responses showed dramatic differences from the observations in previous experiments. Paired *t*-tests were conducted between conditions. For syllables, there was no significant difference between *GP* and *B* in either N1 or P2 response (N1: *t*(16) = −1.052, *p* = 0.308, *d* = −0.182; P2: *t*(16) = −1.068, *p* = 0.301, *d* = −0.12) (Fig. 4*D*). In the *SP*, compared against *B*, the effect was not significant either in the N1 (*SPcon*: *t*(16) = 0.136, *p* = 0.893, *d* = 0.025; *SPinc*: *t*(16) = 1.162, *p* = 0.261, *d* = 0.197) or P2 (*SPcon*: *t*(16) = −0.470, *p* = 0.645, *d* = −0.089; *SPinc*: *t*(16) = −0.662, *p* = 0.517, *d* = - 0.101) (Fig. 4*F*). A repeated-measures one-way ANOVA on responses to tones also did not reveal any significant results (N1: *F*(2,32) = 0.535, *p* = 0.591, partial *η*^*2*^ = 0.032; P2: *F*(2,32) = 0.154, *p* = 0.858, partial *η*^*2*^ = 0.010) (Fig. 4*H*). These null results after attentional manipulation clearly differed from the positive results obtained in motor preparation, suggesting that attention cannot account for the modulation effects observed in Experiments 1 & 2.

### Model simulation results

In a two-layer neural network model (Fig. 5A), we built in the inhibitory and gain modulation functions via a bifurcation of motor signals and an interneuron to simulate the changes of auditory processing across the time course of speech production. During the general preparation (*GP*, left column in Fig. 5B), because there was no speech target, every motor unit activated. The weak activation in each motor unit (dashed line in the first plot) accumulated to a larger activation (bold grey line in the first plot) that induced a strong inhibitory response in the interneuron (solid blue line in the second plot). On the contrary, no motor activation and hence no inhibition in the hearing baseline (*B*, dashed blue line in the second plot). Although the motor activity provided an excitatory gain for each auditory unit in *GP* (solid red line in the second plot) but no gain in hearing (dashed red line at the value of 1 in the second plot), the strong inhibition overwhelmed the excitatory gain and resulted in suppression in auditory responses (the third plot). After temporal averaging of the peak component around 100ms, the simulation results of relative suppression was consistent with our observations in the *GP* condition (the blue bar in the bottom row). The Bayes factor of comparison between the simulation result and results in *GP* condition favored the null (scaled JZS Bayes factor = 3.91), suggesting the model captured the suppressed auditory responses in the *GP*.

During the specific preparation (*SP*, the second column in Fig. 5B), only the motor unit of the prepared target was activated, which yielded the sum of all motor units the same as the unit of the prepared target (in the first plot). This weak motor activation induced a weak inhibition (in the second plot). The relatively stronger gain (in the second plot) that was induced by the specific preparation of the motor target outweighed the inhibition and yielded a response enhancement in the auditory units compared with the hearing baseline (in the third plot). This simulation result of the temporal average was consistent with our observations in the *SP* condition (the red bar in the bottom row). The Bayes factor of comparison between the simulation result and results in *SP* condition favored the null (scaled JZS Bayes factor = 4.21), suggesting the model captures the enhanced auditory responses in the *SP*.

To connect to the previous studies of articulation suppression, we assumed that additional motor neurons in the downstream of speaking action beyond preparation were active (the first plot in the third column). The motor activation could induce a strong inhibition as the auditory suppression occurred near the onset of speech production (Eliades and Wang 2003; Daliri and Max 2016). With the additional inhibitory source, the inhibition to the speech feedback increased (the second plot) and induced a stronger suppression to normal feedback compared to passive listening of playback (the third plot). This model generated a similar result as the speech induced suppression observed in (Houde et al. 2002) (the green bar in the bottom row).

This model can explain the reduction of suppression to the altered feedback (the right column in Fig. 5B). For example, in (Houde et al. 2002) Fig. 7, the suppression of the speech feedback after adding tones becomes much smaller than that to normal feedback. After fixing the parameters in the simulation of articulation induced suppression (similar motor and inhibition in the right column as those in the third column), we fed the auditory input into the auditory unit that was next to the speech target and a unit representing tones to simulate the auditory processing of altered feedback. Because the precise inhibition from articulation to the auditory target reduced its lateral inhibition to its neighbors, the suppression to the altered feedback input became smaller than the suppression to the target (the purple line in the third plot of the right column is higher than the green line in the third plot of the third column). The averaged auditory output was similar to the observation in (Houde et al. 2002) Fig. 7 (the purple bar in the bottom row).

## Discussion

We investigated the functions of motor signals along with the evolution of actions. With a novel delayed articulation paradigm in three electrophysiological experiments, we found that speech preparation at distinct stages differentially modulated auditory neural responses. When no linguistic information was available, the preparatory motor signals ubiquitously suppressed the early neural responses to all speech sounds, whereas the preparatory motor signals generated based on a particular syllable enhanced the neural responses only to the prepared syllable. These modulatory functions in distinct directions along different stages of speech preparation suggest that granular motor signals with different natures were induced along the gradient of action dynamics.

Historically, the terms of *corollary discharge* and *efference copy* were proposed based on the observations of action execution. Arguably, execution is the ending output stage of an action, when presumably all possible motor signals are available. The lack of splitting the potentially complex motor signals may make the inhibitory functions of *CD* overwhelm other functions, yielding the well-established observation of action-induced sensory suppression. However, when considering the processing dynamics and signal contents in the hierarchy of the motor system, distinct motor signals are likely available at different stages (Crapse and Sommer 2008; Straka et al. 2018) and exert distinct modulatory functions on the sensory systems. In this study, the dynamics and contents were experimentally isolated using a delayed articulation paradigm. This experimental manipulation revealed distinct modulatory functions of motor signals along with the evolution of actions, supporting the granular perspective of motor-to-sensory transformation.

Our behavioral and electrophysiological results cumulatively demonstrate that a type of motor signal can be generated during speech preparation even without any preparatory contents. Facilitation in articulation speed was observed after general preparation in both Exp. 1 and 2. Neural suppression in early auditory responses to syllable sounds was also observed in Exp. 2. These behavioral and EEG results were consistent with immense literature about action-induced sensory suppression in both animal models (Poulet and Hedwig 2006; Crapse and Sommer 2008; Eliades and Wang 2008; Schneider et al. 2018; Straka et al. 2018) and humans (Blakemore et al. 1998; Houde et al. 2002). Our results suggest that *CD* provides a uniform inhibitive function that suppresses sensory processing during the action. Moreover, our results reveal that *CD* is a generic form of motor signals that indicate the action, and can be available at the initial stage of action. This is consistent with the function of the *CD* on self-monitoring and agency (Desmurget et al. 2009; Kilteni et al. 2018; Tian et al. 2018).

The *CD* available during the general preparation enhanced the auditory responses to tones. These results suggest that *CD* is generated from and is constrained by the configuration of species’ specific motor system. Although *CD* may not carry any specific content information, it is generated in the motor-to-sensory transformation pathways that adapt to specific actions and reafferent sensory information. In our case, it is human speech — *CD* that is generated from the motor system controlling speech production is sent to and inhibits auditory cortices that represent the human speech sounds. The auditory system that represents pure tones may be relatively spared from inhibition, or its sensitivity maybe even relatively increase because tones are not adapted to human vocal tracks. Therefore, neural systems can separate the ex-afference sensory information (generated from external sources) from re-afference (feedback). These results are also consistent with the previous findings that showed relative increases in auditory responses when the speech feedback is substituted with non-speech sounds (Houde et al. 2002; Christoffels et al. 2011).

Compared with the suppression of speech sounds during general preparation, the motor signals during the preparation of linguistic contents selectively modulated the auditory responses. That is, the motor signals during the specific preparation enhanced the auditory responses to the prepared syllables, whereas induced a mild suppression to unprepared syllables. These results suggest that *efference copy* carries specific content information, and selectively modulates the auditory system that represents the perceptual consequence of speaking. Our results are consistent with recent observations of action induced enhancement (Eliades and Wang 2005, 2008; Flinker et al. 2010; Tian and Poeppel 2013; Tian et al. 2016; Singla et al. 2017; Enikolopov et al. 2018; Cao and Händel 2019; Ma and Tian 2019). Note that the enhancement to the prepared syllables is different from the enhancement to the perturbed feedback (Behroozmand et al. 2009). The observed enhancement during specific preparation is to the prepared syllable. It is more likely caused by the modulation to the speech target, whereas the enhancement to the feedback perturbation occurs during articulation. It fits with the predictive coding in which an error term is induced when the prediction and feedback cannot match.

The distinct directions of modulation effects on sensory processing at different preparation stages offer tantalizing hints suggesting that motor signals of distinct functions are available throughout the entire evolution of action. The *CD* can be available as soon as in the movement intention stage. It dissociates from specific actions that the system will engage, as we observed ubiquitous inhibition in auditory responses to all syllables. Moreover, the *CD* is probably independent of what motor effectors the actions would be executed by, as the auditory suppression was also observed by manual button-press (Bäß et al. 2008; Horváth et al. 2012; Cao et al. 2017). Furthermore, the *CD* is probably general inhibitory motor signals that are available across visual (Sommer and Wurtz 2006), auditory (Poulet and Hedwig 2006), and somatosensory (Blakemore et al. 1998) modalities.

On the contrary, the induction of *EC* requires a concrete action plan. The *EC* that contains specific action information would selectively enhance the perceptual responses to the same information that are contained in the motor signals, as we observed enhancement to the congruent syllable but not incongruent ones in the specific preparation stage. These results agree with the hypothesis of a one-to-one mapping between the motor and sensory systems (Tian and Poeppel 2012, 2013; Tian et al. 2016; Ma and Tian 2019). The enhancement effects of EC reflect the increased sensitivity to the congruent sensory representation, as compared to the incongruent stimuli (Eliades and Wang 2008; Hickok et al. 2011; Ma and Tian 2019). Because the concrete action information can be used to predict detailed perceptual consequences, the function of *EC* would be constrained by the established specific associations between motor and sensory systems, clearly contrasted with the ubiquitous inhibitory function of CD regardless of detailed sensorimotor correspondence.

The observations of dynamics and functional specificity of motor signals inspire upgrades of theories regarding sensorimotor integration and motor control. We first put forward a tentative processing model in the framework of internal forward models with detailed temporal and functional features. *CD* is induced throughout the course of action. It exerts inhibitory functions on given sensory modalities via established motor-to-sensory transformation pathways. *EC* starts later when specific movement parameters are calculated. It increases the sensitivity to a given sensory token that relates to the results of the action via a detailed one-to-one mapping between the motor representation of the specific action and the linked sensory process of perceptual consequences. The complementary functions of *CD* and *EC* collaboratively enable self-monitoring and error detection/correction. The *CD* achieves self-monitoring and agency by suppressing processes in a given sensory modality to indicate the non-specified perceptual consequence of one’s actions. Whereas, the perceptual consequence of an action is sensitized by the *EC* so that the incrementally stronger *CD* when the concrete actions are carried out can precisely inhibit the sensory consequence and indicate possible errors of incorrect sensory feedback.

We quantify the proposed mechanism and potential neural implementation in a computational model (Fig. 5*A*). The copy of motor signals bifurcates. One branch has a direct one-to-one mapping and enhances the postsynaptic gain of the corresponding auditory unit (Ma and Tian 2019). The increase of excitatory gain qualitatively equals to direct excitation but has sustaining effects as those during preparation. Another branch from all motor units activates an interneuron that inhibits all auditory units. During general preparation, activity from all motor units aggregately activates the interneuron that suppresses the neural responses to all syllables (Fig. 5*B*). When detailed information is available in the specific preparation stage, only one motor unit is activated. The excitatory effect from the motor unit outweighs its inhibitory effect and reverses the modulation into enhancement. During action execution, stronger signals from the local motor neurons inhibit the target sound, resulting in speech-induced suppression (Houde et al. 2002). When the auditory feedback is altered, such as tones were mixed into the normal feedback (Houde et al. 2002), the auditory input feeds into the units that are not the speech target. The strong inhibition to the target unit causes less lateral inhibition to the other auditory units and results in a less speech induced suppression to the altered feedback. The simulation results of reduced suppression in off-target units are also consistent with the tuning of suppression observed in the animal models, with less suppression in the responses to the sounds that are farther away from the normal feedback or action-associated sounds (Eliades and Wang 2003; Schneider et al. 2018). That is, a parsimony model of motor signals bifurcation can account for the distinct functions observed in the action preparation and execution stages.

Our model cannot account for the error term that is manifested as a large positive increase in responses to feedback perturbation. For example, the response amplitude of the P2 component increases when the F0 in the feedback is shifted upward (Behroozmand et al. 2009). The response increases are consistent with predictive coding that an error representation can be created by comparing the prediction with the feedback perturbation. However, our simple model only simulates the processes at the basic auditory levels. The representation of comparison results (the increases) is usually modeled at an upper level (Arnal and Giraud 2012), which is consistent with the common observations of response increases at a later latency, for example, P2 but not P1 (Behroozmand et al. 2009). The lack of such computation hierarchy in our simple model makes it hard to explain the error increases, but rather to concentrate on explaining the direct modulations on auditory processing.

The model is inspired by our novel empirical findings of distinct modulation effects of speech preparation. This model is a quantification of a possible algorithm and neural implementation for the observed functional distinctions between *efference copy* and *corollary discharge*. It offers a novel perspective to approach the motor-to-sensory transformation and internal forward models. It also provides testable hypotheses regarding the relations between motor and perceptual processes, as well as the computations and neural implementations that mediate the interaction among systems, such as the gain modulation of efference copy and its neural implementations, the extent of the neural representation of motor units and their weak activation during general preparation, the connectivity of interneurons and the extent of their inhibitory functions on different types of sensory representations, the stronger and precise inhibition of motor signals near and during action execution.

The proposed mechanism can be tested in different sensory modalities in both human and animal models. For example, the different timing and weighting of *CD* and *EC* could be realized by the onset of motor signals from different cortical areas in the motor hierarchy. In the visual domain, it could be the difference between the up-stream LIP for *CD* and downstream FEF for *EC* (Zirnsak et al. 2014; Wang et al. 2016). In the auditory domain, it could be the intention to speak in IPS (Tian and Poeppel 2010) for the initialization of *CD* and frontal motor regions (including pre-motor, SMA, IFG) for *EC* (Tian et al. 2016). Moreover, motor signals are theorized to convey predictive signals to facilitate auditory perception and auditory-guided behaviors (Schneider et al. 2014). The different functions of motor signals could be manifested in the plasticity and modulation. In vision, the stabilization (temporal inhibition of visual processing) during saccades, remapping of the receptive field before saccades, and partial active receptive field during saccades (Sommer and Wurtz 2006; Wang et al. 2016) could be caused by the interplay of distinct motor functions that modulate the visual processing. In audition, learning, self-monitoring of own articulation, differential manipulation of sensitive to auditory target, and speech error detection and correction (Houde 1998; Hickok et al. 2011; Hickok 2012; Tian and Poeppel 2014; Liu and Tian 2018) could be mediated by the interaction of distinct motor functions that modulate the auditory processing. The specific functions and neural pathways about the proposed distinct motor signals could be further investigated and mapped out by electrophysiological, neuroimaging, and optic-genetic approaches (Poulet and Hedwig 2006; Sommer and Wurtz 2006; Schneider et al. 2014, 2018).

Our results may offer insights about the cognitive neural mechanisms that mediated clinical and mental disorders. For example, our results may implicate a possible cause of auditory hallucinations from a perspective of internal monitoring and control. The normal population may use *EC* to internally induce auditory mental images and use the inhibitory function of *CD* to ‘label’ the source as internally self-generated. This interplay between *CD* and *EC* separates mental imagery from reality. However, patients suffering from auditory hallucinations may have intact *EC* to generate auditory mental images internally, whereas the inhibitory *CD* malfunctions (Tian and Poeppel 2012; Yang et al. 2019). The intact enhancement function of *EC* generates auditory and speech representation based on internal stimulation of motor signals, but the lack of suppressive function of *CD* fails to label the internally generated sounds as self-generated. The internal prediction of a perceptual consequence, which has the same neural representation as an external perception, is erroneously interpreted as the result of external sources, which results in auditory hallucinations. Results in the current study support the hypothesis that two distinct motor signals are available to modulate perceptual responses, indicating their possible roles in speech monitoring and control, as well as the potential causes of auditory hallucinations.

Using a novel delayed articulation paradigm, we observed that distinct motor signals were generated in the motor-to-sensory transformation and integrated with sensory input to modulate perception during speech preparation. The content in the motor signals available at distinct stages of speech preparation determined the nature of signals — *corollary discharge* or *efference copy* and constrained their modulatory functions on auditory processing.

## Acknowledgment

We thank Xingye Chen for her help in running experiments. This study was supported by the National Natural Science Foundation of China 31871131, the Major Program of Science and Technology Commission of Shanghai Municipality (STCSM) 17JC1404104, and the Program of Introducing Talents of Discipline to Universities, Base B16018.

